# A novel approach to increase glial cell populations in brain microphysiological systems

**DOI:** 10.1101/2023.09.14.557775

**Authors:** Itzy E. Morales Pantoja, Lixuan Ding, Paulo E. C. Leite, Suelen A. Marques, July Carolina Romero, Dowlette-Mary Alam El Din, Donald J. Zack, Xitiz Chamling, Lena Smirnova

## Abstract

Brain microphysiological systems (bMPS), which recapitulate human brain cellular architecture and functionality more closely than traditional monolayer cultures, have become a practical, non-invasive, and increasingly relevant platform for the study of neurological function in health and disease. These models include 3D spheroids and organoids as well as organ-on-chip models. Currently, however, existing 3D brain models vary in reflecting the relative populations of the different cell types present in the human brain. Most of the models consist mainly of neurons, while glial cells represent a smaller portion of the cell populations. Here, by means of a chemically defined glial-enriched medium (GEM), we present an improved method to expand the population of astrocytes and oligodendrocytes without compromising neuronal differentiation in bMPS. An important finding is that astrocytes not only increased in number but also changed in morphology when cultured in GEM, more closely recapitulating primary culture astrocytes. We demonstrate oligodendrocyte and astrocyte enrichment in GEM bMPS using a variety of complementary methods. We found that GEM bMPS are electro-chemically active and showed different patterns of Ca^+2^ staining and flux. Synaptic vesicles and terminals observed by electron microscopy were also present. No significant changes in neuronal differentiation were observed by gene expression, however, GEM enhanced neurite outgrowth and cell migration, and differentially modulated neuronal maturation in two different iPSC lines. Our results have the potential to significantly improve in vivo-like functionality of bMPS for the study of neurological diseases and drug discovery, contributing to the unmet need for safe human models.

## 1. Introduction

Brain MPS provide an accessible and relevant platform for studying states of homeostasis and disease. However, most brain organoid models are predominantly composed of different neuronal types, with a smaller percentage of glial cells present than exists naturally (1–3), and so improving a cellular composition to be more naturally reflective of the physiological cellular composition of the brain is crucial to advancing 3D *in vitro* human systems.

The exquisite functionality of the human brain is derived in part from the complex interplay between glial and neuronal cells. While the term glial encompasses microglia, astrocytes, and oligodendrocytes, the focus of the present study is on astrocytes and oligodendrocytes. Microglia can be differentiated separately and incorporated into the organoid cultures (4–6).

Glial cells are essential for the proper development and function of the central nervous system (CNS)(7) even though historically they have been regarded as neuron-supporting cells (8). Some of the supportive yet fundamental roles of astrocytes are to provide mitochondria and neurotransmitter components to neurons and regulate glutamate uptake (9,10).

Oligodendrocytes provide the myelin sheath that insulates axons and allows efficient conduction of electrical signals in the CNS and provide trophic support to the axons, which is crucial for axonal integrity (11). A robust body of research has demonstrated that astrocytes and oligodendrocytes have multifaceted roles in synapse formation and pruning, and both have active roles in building and maintaining neuronal networks, directly impacting plasticity and cognitive function (8,9,12–14). These paramount roles shifted the dogma away from the notion that neuron-to-neuron interactions were unique to brain complexity under homeostasis and disease and towards a new paradigm in which the importance of glia-to-neuron interplay was recognized. Both astrocytes and oligodendrocytes are intrinsically involved in the formation and modulation of brain function (15). Recent studies in adult mice have shown that oligodendrocytes are necessary for motor learning, and that differentiation of OPCs into oligodendrocytes occurs at the early stages of the learning process along with synaptic changes (16). Astrocytes induce long-term potentiation and promote memory allocation (17,18), as they are able to provide signals for the formation of new neurons and facilitate their integration into established circuits (13). Both oligodendrocytes and astrocytes modulate the formation of short, recent, and remote memory (15), and their absence is associated with disorganized white matter architecture, loss of blood–brain barrier integrity, dysmyelination, cognitive impairment and behavioral dysfunction (9,19,20).

Despite many advances, much remains unknown about the normal and pathologic progression of events in the adult brain. The inaccessibility of the cells of interest and the lengthy timeframe over which events happen are restrictive, but recent advancements in stem cell technology have enabled new avenues of study. Further, MPS, which closer recapitulate organ functionality, are allowing for steady improvements of *in vitro* human models. Thus, increasing the oligodendrocyte and astrocyte populations in 3D brain models offer the potential to better understand human neurobiology in states of homeostasis and disease.

The ability to study glial cells in an environment that more closely resembles a native cellular composition is important, as it has been demonstrated that primary astrocytes change both their form and function after isolation from the brain tissue (21). In this work we significantly increased the glial population in bMPS in a relatively short time, bringing the model closer toward physiological interactions between neurons and glial cells.

## 2. Materials and Methods

### 2.1 Differentiation of brain MPS from pluripotent stem cells

Brain MPS were differentiated from two induced Pluripotent Stem Cell (iPSC) lines; NIBSC-8 (female origin, UK National Institute for Biological Standards and Control (NIBSC)), and CS2PFYiCTR-nx.x (here CS2, male origin from Cedars-Sinai Medical Center Stem Cell repository), and from two embryonic stem cell (ESC) lines, H9 and Reus1, which were previously genetically modified (22). All cell lines are mycoplasma-free, with normal karyotypes, and were differentiated following our previously published two-step protocol with modifications to reflect glia enrichment conditions (Romero et al., 2023). Briefly, hiPSCs and hESCs were cultured in mTESR-Plus medium (StemCell Technologies at 5% O_2_, 5% CO_2_ and 37 °C). The cells were characterized based on expression of stemness markers (Oct4, Nanog and TRA-1-60) by flow cytometry (iPSC characterization see Romero et al., 2023(23), ESC characterization is summarized in Supplemental Figures 1 and 2). Stem cells were then differentiated in a monolayer to neuroprogenitor cells (NPCs) using a serum-free, chemically defined neural induction medium (Gibco, Thermo Fisher Scientific). NPCs were expanded and quality controlled for expression of NPS markers, Nestin, Sox2 (Romero et al., 2023 (23) and Supplemental Figures 3 and 4). To initiate bMPS differentiation, a single cell suspension of 2 × 10^6^ NPCs was distributed per well into uncoated 6-well plates. Cultures were kept at 37°C, 5% CO_2_, and 20% O_2_ under constant gyratory shaking (80 rpm, 19 mm orbit) to form 3D aggregates. After 48 hours, differentiation was induced with serum-free, chemically defined standard differentiation medium (SDM): B-27™ Electrophysiology kit, 2% Glutamax (Gibco, Thermo Fisher Scientific), 10 ng/ml human recombinant Glial Cell Derived Neurotrophic Factor (GDNF) (GeminiBio™), 10 ng/ml human recombinant (Brain Derived Neurotrophic Factor (BDNF) (GeminiBio™), 1% Pen/Strep (Gibco, Thermo Fisher Scientific). Approximately 75% of the medium was changed three times a week. Standard differentiation medium (SDM) was used as the standard/control culture conditions throughout the entire differentiation process, and it was the base for glia-enriched media (GEM). For GEM conditions from week 2 to 8 of differentiation, SDM was supplemented with 10ng/mL PDGF-AA, 10 ng/mL IGF-1, 60 ng/mL T3, 100 ng/mL biotin, 1 µM cAMP, and 25 µg/mL insulin. For terminal differentiation from week 8 to week 15 in GEM conditions (GEMtd), SDM was supplemented with 60 ng/mL T3, 100 ng/mL biotin, 1 µM cAMP, 25 µg/mL insulin, 20 µg/mL ascorbic acid, and 10 mM HEPES.

### 2.2 Immunofluorescence

BMPS were fixed in 4% paraformaldehyde for 45 min at 4°C, washed three times in 1X PBS, and permeabilized with 1 ml permeabilization solution (1X PBS, 0.1% Triton X) for 30 min at 4°C. Cultures were transferred to a 24-well plate and blocked with 100% BlockAid™ (Thermo Fisher Scientific) on a shaker for 1 h at 4°C. Then bMPS were incubated at 4°C for 48 h with a combination of primary antibodies (Table 1) in 10% BlockAid™, 1% BSA (Bovine Serum Albumin) and 0.1% Triton X in PBS. bMPS were washed in PBS three times and incubated with secondary antibody for 24 h in 10% BlockAid™, 1% BSA and 0.1% TritonX in PBS. After three 30 min washes, bMPS were incubated overnight with goat anti-mouse secondary antibody conjugated with Alexa 488, or Alexa 647 (1:500, Thermo Fisher Scientific) and Hoechst 33342 trihydrochloride (1:2000, Invitrogen). bMPS were mounted on slides with coverslips and Prolong Gold antifade reagent (Molecular Probes); negative controls were processed omitting the primary antibodies. Images were taken with a Zeiss UV-LSM 700 confocal microscope. Further imaging processing and quantification were performed using ImageJ2 version 2.9.0/1.53t.

**Table 1.** Primary antibodies used for immunofluorescence.

| Antibody | Host | Type | Source / Cat No. | Dilution |
| --- | --- | --- | --- | --- |
| GFAP | Rabbit | Polyclonal | DAKO Z0332 | 1:400 |
| Nestin | Rabbit | Polyclonal | Sigma N5413-100UG | 1:200 |
| NF-H | Chicken | Polyclonal | Invitrogen PA1-10002 | 1:1000 |
| O4 | Mouse | Monoclonal | R&D Systems MAB1326 | 1:200 |
| Sox 2 | Mouse | Monoclonal | Invitrogen MA1-014 | 1:200 |
| β-Tubulin Class III | Mouse | Monoclonal | Sigma T5076 | 1:1500 |

### 2.3 Flow cytometry analysis

90–100% confluent ESC and NPC were washed with PBS and treated with gentle cell dissociation reagents (GCDR) for 5 min at RT. After full dissociation, 1 ml of Hibernate E medium was added, and the cell suspension was collected into flow cytometry tubes (Corning™Falcon™) and spun down at 300 × *g* for 4 min at RT. The pellet was resuspended in 1 ml of 1X PBS and 1 μl of Fixable Viability Stain 780 (BD Horizon) and incubated for 5 min at 37°C protected from light. After incubation, 2 ml of Stain Buffer (BD Biosciences) was added, and cells were centrifuged at 300 × *g* for 4 min at RT. Pellet was then resuspended in 500 μl of 2% PFA (Sigma Aldrich) and cells were fixed for 20 min at 4°C. A total of 500 μl of permeabilization solution [1X PBS, 1% Triton™ X-100 (Sigma-Aldrich)] were then added for 10 min at RT. A total of 2 ml of washing solution 1 (WS1: 1X PBS, 1% (BSA, Sigma), 0.1% Triton X) was then added to the cells and centrifuged at 300 × *g* for 4 min at 4°C. The cells pellet was resuspended in 300 μl of blocking solution (1X PBS, 1% BSA, 10% normal goat serum, 0.1% Triton™ X-100) and incubated for 1 h at 4°C. A total of 1 ml of WS1 was added and samples were centrifuged at 300 × *g* for 4 min at 4°C. The cells were then resuspended in 100 μl of blocking solution and conjugated antibodies (Table 2) were added directly to each tube and incubated for 1 h at 4°C in the dark. Next, 2 ml of WS1 was added to each sample and cells were centrifuged at 300 × *g* for 4 min at 4°C. The wash step was then repeated once. For the third wash, 2 ml of washing solution 2 (WS2: 1X PBS, 1% BSA) was used. The cells were then resuspended in 300 μl of WS2 and analyzed on the flow cytometer (SONY SH800). For compensation and gating, single-color controls were included for each experiment. Unstained cells were used as a control for viability, and compensation beads were used for antibody staining [anti-mouse Ig, k/negative control Compensation Particles set, BD Biosciences, and UltraComp eBeads™ Compensation Beads (Thermo Fisher Scientific)]. Compensation was performed before running the samples, and then later when processing the results. Flow cytometry data was processed and visualized using FlowJo 10.8.1 (Becton Dickinson & Company).

**Table 2.** Primary antibodies used for flowcytometry.

| Antibody | Host | Type | Source / Cat No. | Dilution |
| --- | --- | --- | --- | --- |
| Oct 4 | Mouse | Monoclonal | BD Bioscience 560253 | 1:5 |
| Nanog | Mouse | Monoclonal | BD Bioscience 561300 | 1:20 |
| Nestin | Mouse | Monoclonal | BD Bioscience 560396 | 1:20 |
| Sox 2 | Mouse | Monoclonal | BD Bioscience 561506 | 1:20 |
| Tra-1-60 | Mouse | Monoclonal | BD Bioscience 561573 | 1:20 |

### 2.4 RT-qPCR

Total RNA was isolated via a Quick-RNA™ Microprep Kit (Zymo Research). RNA quantity and purity were determined using NanoDrop™ 2000c Spectrophotometer (Thermo Fisher Scientific). A total of 500 ng of RNA were reverse-transcribed using M-MLV Reverse Transcriptase and Random Hexamer primers (Promega) according to the manufacturer’s instructions. The expression of genes was evaluated using the TaqMan gene expression assay (Applied Biosystems) (Table 3). Real-time quantitative PCR (RT-qPCR) was performed using a 7500 Fast Real-Time system machine (Applied Biosystems). Gene expression was normalized to the expression of the reference gene Actin beta (ACTB).

**Table 3.** Primers used for RT-qPCR analysis.

| Assay name | Assay ID | Assay type | Catalog number |
| --- | --- | --- | --- |
| ACTB | Hs01060665 | TaqMan® Gene Expression Assay | 4331182 |
| GFAP | Hs00909233 | TaqMan® Gene Expression Assay | 4331182 |
| MBP | Hs00921945 | TaqMan® Gene Expression Assay | 4331182 |
| RBFOX3 (NeuN) | Hs01370653 | TaqMan® Gene Expression Assay | 4331182 |

### 2.5 Bioluminescence Nanoluc assay

Nanoluc activity was measured using NanoGlo luciferase assay reagents (Promega N1150) following manufacture’s instruction. Briefly, 50uL of NanoGlo reaction mix was added to each well containing 50 µL of medium. RLU reading was performed with a microplate reader (Promega Glomax Multi Detection) using the luminescence protocol.

### 2.6 Neurite Outgrowth assay

Ten bMPS per well were seeded on black, 24-well glass-bottomed plates coated with Poly-L-ornithine/laminin (15 µg/mL Sigma Aldrich 10 µg/mL Sigma Aldrich), respectively. Organoids were incubated at 37°C for five days statically. After five days, the bMPS were fixed with 4% PFA for 45 min at 4°C, followed by three rinses with 1% BSA in PBS (washing solution 1, WS1), permeabilized and blocked for 1h in 10% goat serum, 1% BSA, and 0.15 % saponin in PBS (blocking solution). BMPS were incubated at 4°C for 24 h with antibodies against β-III-tubulin and GFAP (Table 1) in blocking solution. bMPS were then washed in washing solution 2 (WS2), composed of 1% BSA and 0.15 % saponin three times and incubated with nuclear dye Hoechst and secondary antibodies for 24 h in blocking solution at 4°C. bMPS were washed three times with WS1 and imaged with an ECHO laboratory Revolve microscope with 4/0.13 magnification objective. Neurite density and length were quantified using the ImageJ Sholl plug-in (24). The ratio was calculated for each shell (number of intersections/distances from the edge of the organoid) and plotted. The area under the curve (AUC) was calculated for each organoid and then averaged.

### 2.7 Calcium imaging

Brain MPS were incubated with 10 µM Fluo-4AM (Tocris) and 0.5% Pluronic™ F-127 (Thermo Fisher Scientific) acid in the dark for 2 h at 37°C and 5% CO_2_ without shaking. After three washing steps, organoids were transferred to a glass-bottomed 24-well plate to be imaged with the resonant scanner of an Olympus FV3000-RS with a speed of 12.5 frames per second for six minutes. After time series images were taken for all conditions, Z-stacks of 20 µm were taken. bMPS were kept at 37°C and 5% CO2 throughout imaging. Raw fluorescent intensities of whole bMPS over time were analyzed using the ROI manager in ImageJ. These raw fluorescence intensities were further analyzed using Excel to obtain delta F/F plots. Additionally, Z-stacks collapsed into Z-projections using ImageJ.

### 2.8 Transmission Electron Microscopy

The bMPS were fixed with 4% paraformaldehyde and 2.5% glutaraldehyde in sodium cacodylate buffer (pH7.4), and then washed for 10 minutes in the same buffer three times. The bMPS were post-fixed in a 1% osmium tetroxide (OsO4) solution plus 0.8% potassium ferrocyanide and 5 nM calcium chloride in sodium cacodylate buffer. Brain MPS were then washed with cacodylate buffer and distilled water and incubated in 1% uranyl acetate overnight. After gradual dehydration with acetone series (30, 50, 70, 90, and 100%), bMPS were embedded in Poly/Bed 812 resin (Ted Pella Inc, USA) and polymerized at 60°C for 48 h. The ultrathin cross-sections (70 nm) were obtained with an RMC ultramicrotome. Transmission electron microscope (TEM) analyses were performed using the TEM (JEM 1011, Jeol) operated at 80 kV.

### 2.9 Statistical analysis

All graphs and statistical analysis were done in GraphPad Prism, version 9. 5.1. Statistical methods used are specified in each figure. Grouped analysis was performed by Mann-Whitney test for figures 2, 3 and 4C. Gene expression time course was analyzed using two-way ANOVA test with Holm-Šidák’s post-hoc test for multiple comparison for figure 4A-B. Student unpaired t-test was used for significance for figure 5, and two-way ANOVA test with Šidák’s multiple comparisons for supplemental figure 6. P < 0.05 was considered significant.

## 3. Results

### 3.1 GEM increases astrocyte and oligodendrocyte populations and differentially modulates neuronal maturation

During the first two weeks of bMPS differentiation, standard bMPS differentiation medium was used (25). In the previously established standard protocol developed by Pamies et al., 2017 (25), BDNF and GDNF drive the 8-week neural differentiation. In the present method, additional key factors were added to enhance glial differentiation and promote cell survival: T3 (3,3,5-Triiodo-l-thyronine) (26,27), IGF-1 (Insulin like growth factor 1) and PDGF-AA (Platelet-derived growth factor AA). IGF-1 is one of the primary factors regulating cell survival, proliferation, and maturation while PDGF-AA is involved in oligodendrocyte progenitor cells (OPCs) migration (28–30). Studies utilizing fetal and adult brain dissociated tissue to investigate the response of growth factors have shown that IGF-1 increased the expression of GC (galactocerebroside), and PDGF-AA increased expression of O4/GC-positive cells (31). A combination of BDNF and PDGF-AA promote neural cell differentiation towards both neuronal and oligo lineages (32), whereas PDGF-AA specifically enhances oligo commitment (28,33).

The glia enrichment regimen was initiated at week two of differentiation to expand and (re)direct the existing pool of NPCs and OPCs towards glial lineage commitment (Figure 1). Thus, from the second to eighth week, bMPS were cultured either in Glia Enrichment Medium (GEM) or Standard Differentiation Medium (SDM). From the eighth to tenth week, the GEM cultures were switched to GEMtd (<u>t</u>erminal <u>d</u>ifferentiation), while the control bMPS remained in the same standard differentiation medium throughout the entire differentiation process.

**Figure 1.**
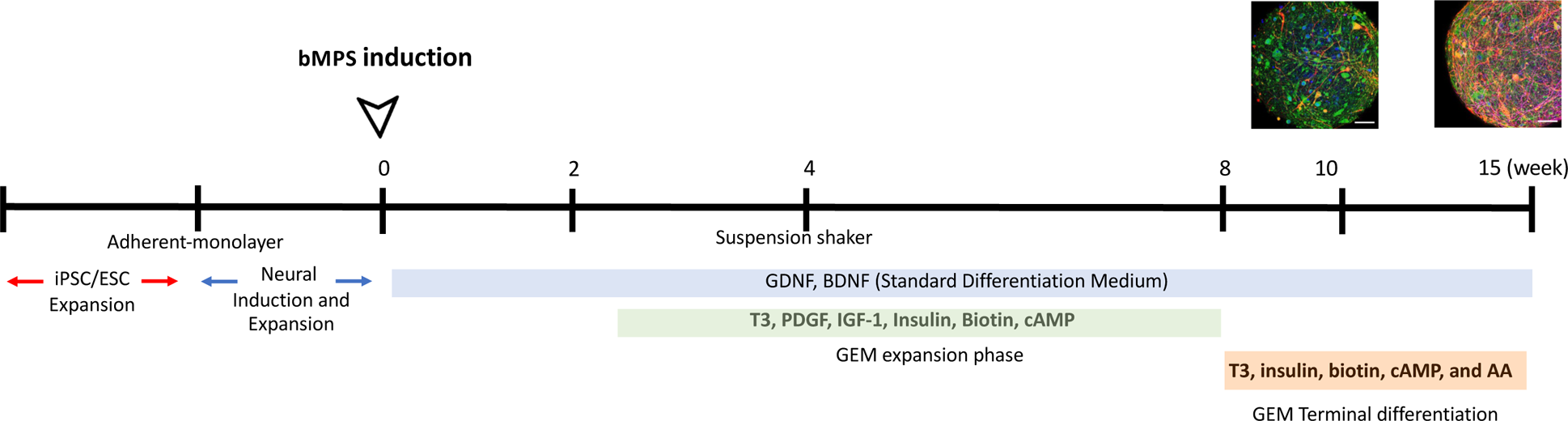
Timeline of brain organoid differentiation. Schematic representation of the timeline of differentiation, GEM phases, and GEM key components.

To characterize the effects of GEM, bMPS were immunostained for canonical glial and neuronal markers at weeks 10 and 15 of differentiation. We found that GEM increased the presence of O4+ oligodendrocytes and GFAP+ astrocytes in comparison to SDM at both time points. The effect was reproducible in two iPSC lines used, namely NIBSC8 and CS2 (Figures 2 and 3). O4 expression and morphology changed as the bMPS matured, and the effect was particularly stronger in GEM-treated cultures. GEM-O4+ cells displayed longer processes at week 15 (Figures 2A and 3A). Similar effects were also observed for GFAP+ cells, where both time in culture and medium composition of GEM significantly increased GFAP+ astrocytes in the bMPS. GEM-astrocyte morphology better recapitulated the morphology of primary cultures and appeared to induce a fibrous phenotype (34) (Figures 2B, 3B, and Supplemental Figure 5).

**Figure 2.**
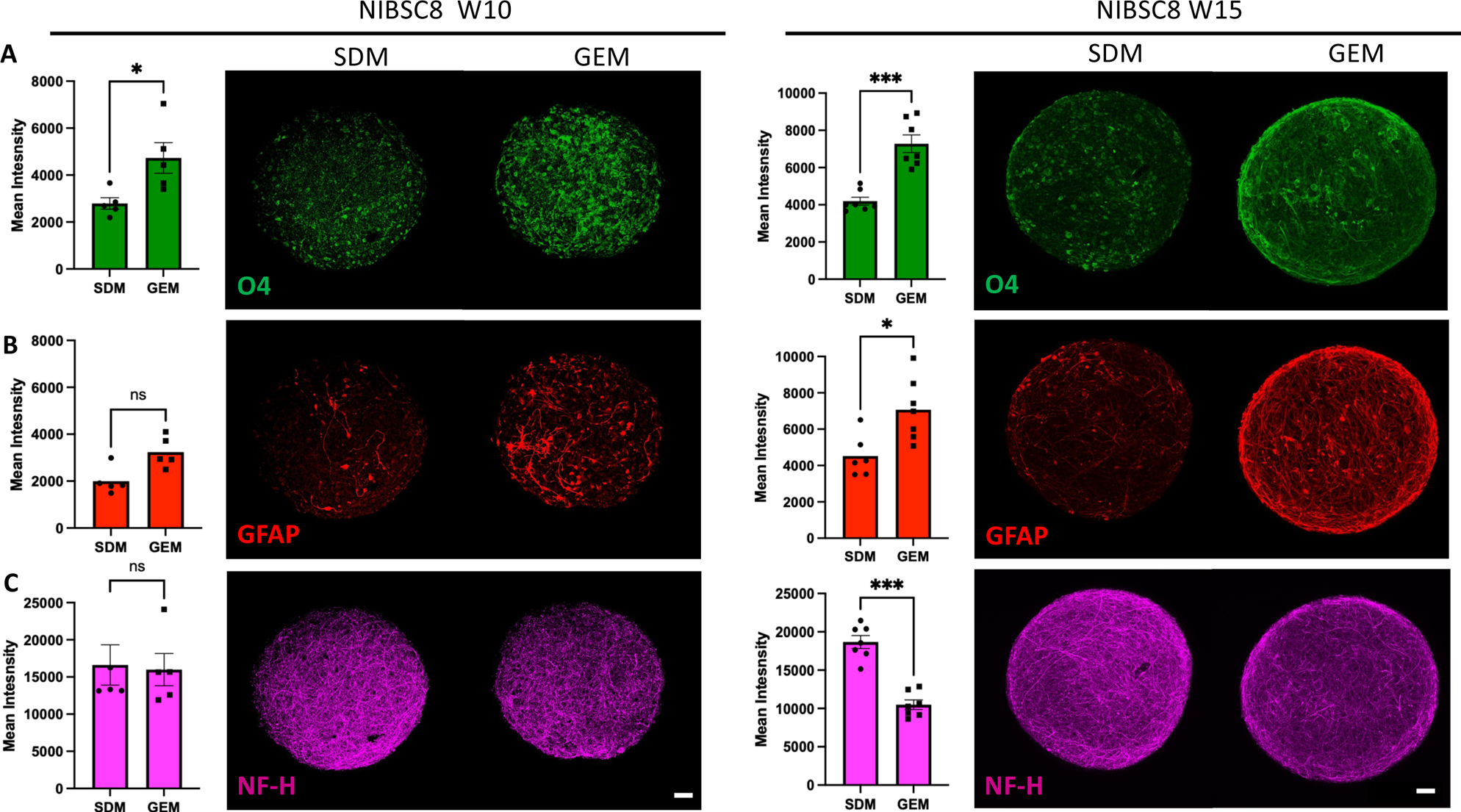
Figure 2. Quantification of glial and neuronal markers in NIBSC8 iPSC line differentiated to bMPS in SDM and GEM for 10 and 15 weeks. Data represents mean intensity ± SEM of O4- (A, green), GFAP- (B, red), and NF-H- (C, purple) positive cell from 5-7 bMPS per condition and time point obtained from 8-11 stacks from z-stacks of one-micron intervals. Mann-Whitney test was used for significance testing. Representative images from at least 3 experiments are shown. The scale bar represents 50 µm.

**Figure 3.**
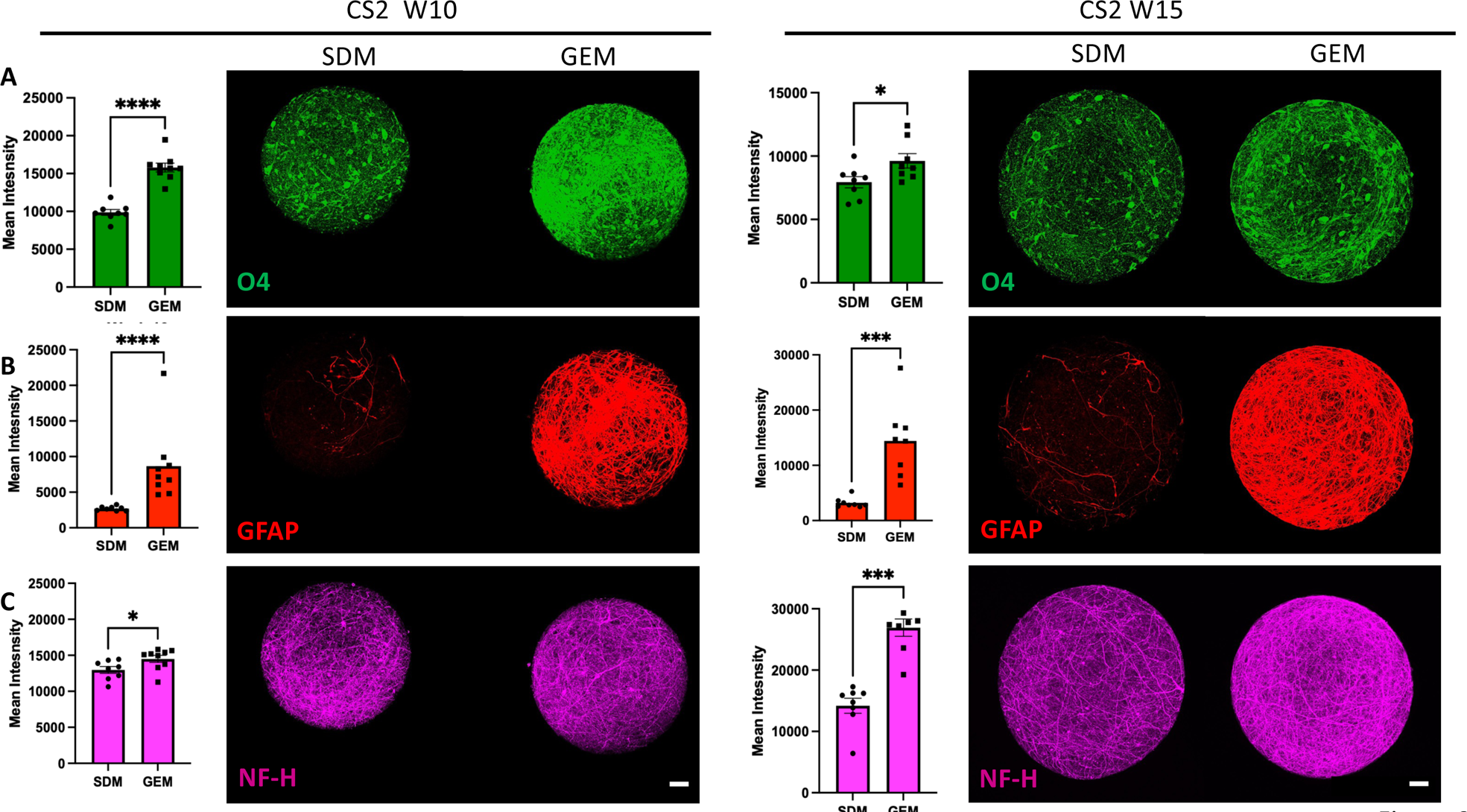
Quantification of glial and neuronal markers in CS2 iPSC line differentiated to bMPS in SDM and GEM for 10 and 15 weeks. Data represents mean intensity ± SEM of O4- (A, green), GFAP- (B, red), and NF-H- (C, purple) positive cell from 5-7 bMPS per condition and time point obtained from 8-11 stacks from z-stacks of one-micron intervals. Mann-Whitney test was used for significance testing. Representative images from at least 3 experiments are shown. The scale bar represents 50 µm.

NIBSC8 line showed no effect on the expression of the neuronal marker NF-H at week 10, however, the mean intensity of NF-H significantly decreased at week 15 (Fig. 2C), while CS2-line showed an increase of NF-H intensity at both time points (Fig. 3C).

### 3.2 GEM dramatically increases the gene expression of *GFAP* and enhances the expression of *MBP* in bMPS from two iPSC and two ESC lines respectively

We quantified the relative gene expression of *MBP, GFAP*, and *NeuN* to further confirm the differences in the presence of astrocytes, oligodendrocytes, and neurons in GEM versus SDM. We found similar patterns of expression in both iPSC lines, albeit with minor variability in the timing of enriched expression. Although differences in *MBP* expression did not reach statistical significance, at 10 weeks, *MBP* expression in GEM organoids was 1.7 times higher than in SDM in NIBSC8 (Figure 4A) and 1.2 times higher in CS2 (Figure 4B). *GFAP* expression in GEM vs. SDM organoids significantly increased by 64 times and 37 times at week 10 in NIBSC8 and CS2, respectively (Figure 4A and B) and there was no difference between SDM and GEM organoids in the relative expression of the postmitotic neuronal marker *NeuN* at any time point of differentiation in either cell line (Figure 4A and B).

**Figure 4.**
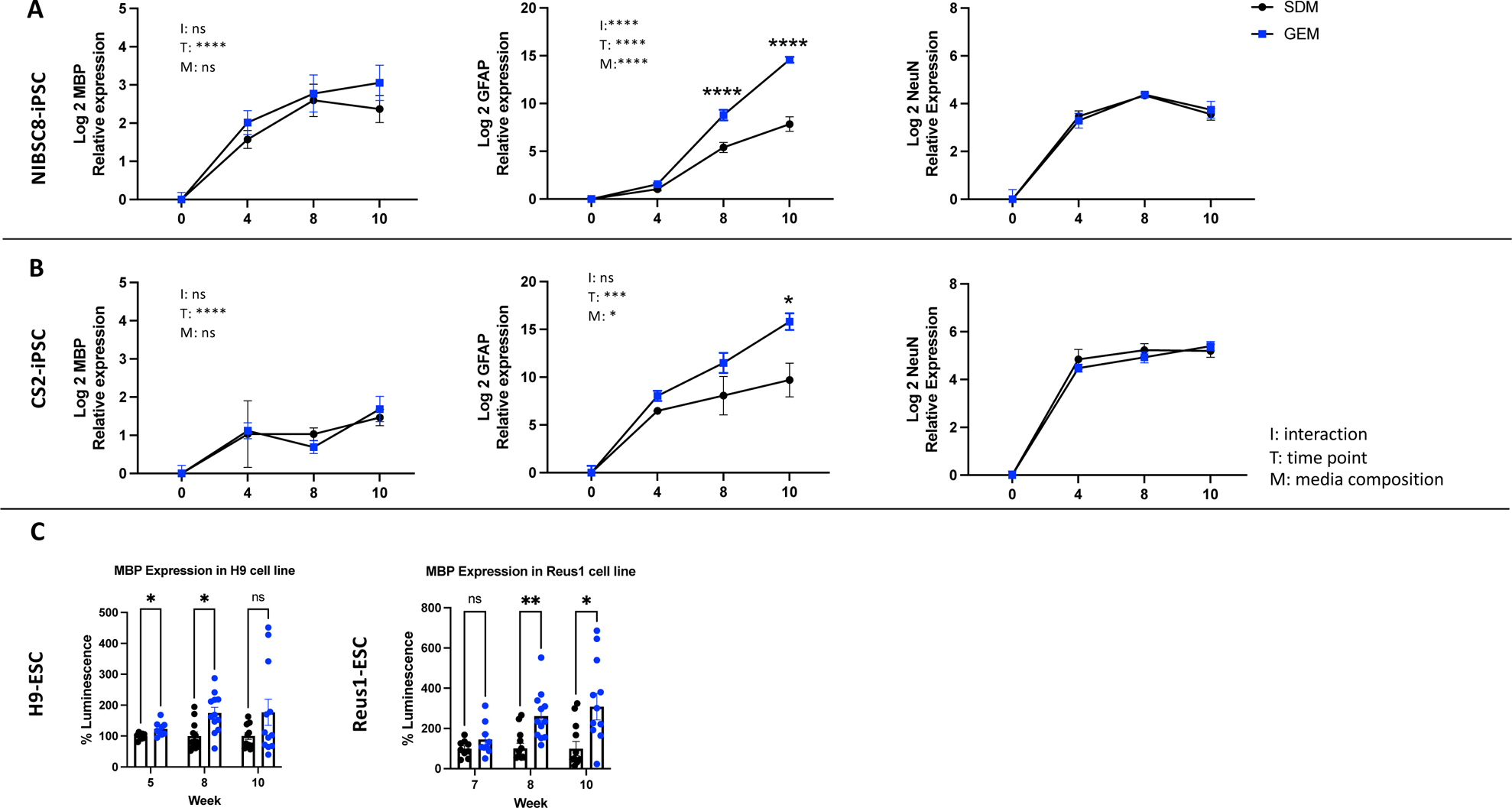
Gene expression of glial and neuronal markers in GEM and SDM. Relative expressions of *MBP* (oligodendrocyte marker), *GFAP* (astrocyte marker), and *NeuN* (post mitotic neuronal marker) are shown for (A) NIBSC8 and (B) CS2 cell line. Data represents mean ± SEM from 3-6 independent experiments per time point, per condition, per cell line. Two-way ANOVA with Holm-Sidak post-hoc test was used for significance test, where “I” stand for interaction, “T” for time point, and “M” for medium composition. (C): Expression of MBP, quantified by luciferase assay in two MBP-nanoluc reporter lines (Reus1 and H9). Data represents mean ± SEM from two independent experiments with total of 6 bMBP for weeks 5 and 7 and 12 bMPS for weeks 8 and 10 measured per cell line.

To confidently establish oligodendrocyte enrichment, we employed Reus1 and H9 ESC lines engineered with CRISPR-Cas9-based knock-in Nanoluc (secNluc) reporter driven by the myelin basic protein (MBP) promoter. In this model, secNanoLuc is separated from the *MBP* gene product by a 2A sequence (22). NanoLuc is secreted into the culture media when MBP is expressed and allows quantification of MBP promotor activity as proxy for MBP gene expression using a small aliquot of the cell culture media in a quick and reliable manner. GEM significantly increased the expression of MBP in both ESC lines at 8 weeks of differentiation. At 10 weeks, MBP levels were higher in both cell lines, but due to higher variability in H9 cells the significance was not reached although a clear trend was observed (Figure 4C).

### 3.3 GEM enhances neurite outgrowth and glial migration in bMPS derived from two iPSC lines

Neurite outgrowth is an important process for the connectivity and wiring of the CNS and necessary for regeneration (35). Astrocytes play an integral role regulating this process by providing molecular cues and serving as a supporting scaffold (36). Given the importance of this process, in addition to the characterization of canonical markers, we sought to investigate the ability of the cultures to develop neurites. For this functional assay, mature 12 to 13 week-bMPS were switched from suspension and constant gyratory shaking to static culture conditions, allowing the organoids to adhere to a laminin extracellular matrix for four days.

GEM-treated cultures exhibited significantly greater neurite length, thickness, and density, as well as astrocyte migration (Figure 5A-D). We observed a clear structural alignment of neurites and glial processes, as well as the soma of neurons and astrocytes away from the organoid, indicating successful cell migration. While GEM significantly enhanced neurite outgrowth and cell migration in both cell lines, the effect was stronger in NIBSC8 line (Figure 5A, C).

**Figure 5.**
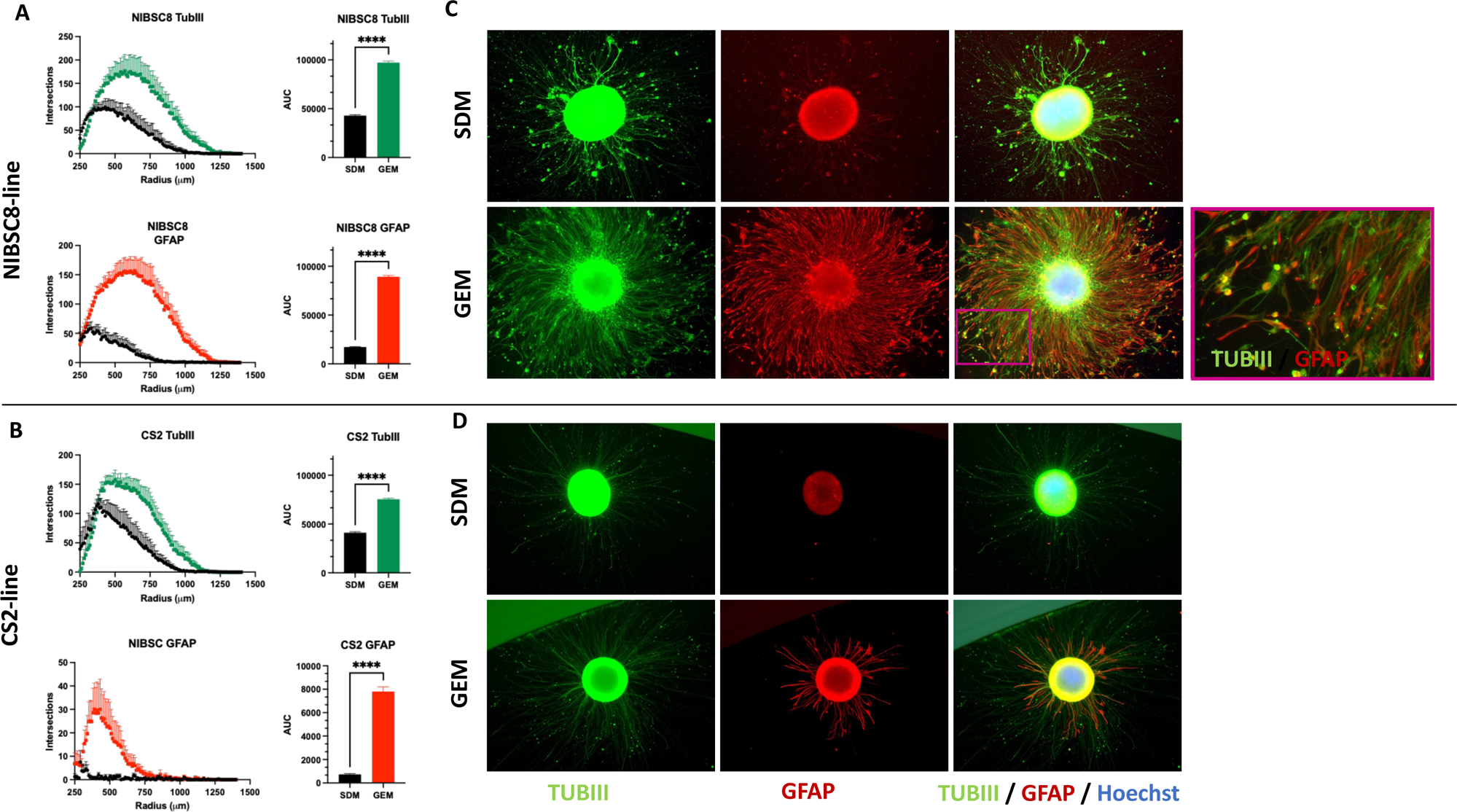
Neurite outgrowth and astrocyte migration in bMPS cultured in SDM and GEM. Sholl analysis of differences in neurite outgrowth (top) and astrocyte migration (bottom) in NIBSC8-line (A) and CS2 (B) between SDM and GEM conditions showing number of intersections depending on the distance from the edge of the organoid. The area under the curve (AUC) was calculated for each condition, and each graph represents the AUC mean ± SEM from 5 organoids. Unpaired t-test was used to calculate significance. Representative images of organoids derived from NIBSC8 (C) and CS2 (D) from two independent experiments and ten bMPS screened per condition. Neurons are labeled with antibody against beta-III-tubulin (TUB III, green) and astrocytes are stained with GFAP antibody (GFAP, red). Nuclei are stained with Hoechst (blue). 5x magnification.

### 3.4 Visualization of myelination, synaptic vesicles (SV), and terminals (ST) in bMPS

Rapid neuronal communication requires proper calcium influx response that triggers the fusion of synaptic vesicles with presynaptic terminals, i.e., synaptic transmission (37). Myelination is evidence of mature and functional oligodendrocytes, and is also required for proper electrical signal propagation along neurons. We assessed the synaptic terminals and myelination at the ultrastructural level by TEM at weeks 8 and 13 of differentiation, as these time points denote the transition from glial-enrichment expansion to terminal differentiation and maturity. bMPS cultured in GEM showed a variety of cellular profiles with a more organized ultrastructure and displayed microtubule-rich processes (e.g. Figure 6A c,d yellow arrow) indicative of incipient neuronal differentiation, and a high density of synaptic vesicles and synaptic terminals in CS2, H9, and Reus1 cell lines (Figure 6Ae, 6Af, 6Db, and 6Gd). We observed myelination in all four cell lines and in all conditions and time points.

**Figure 6.**
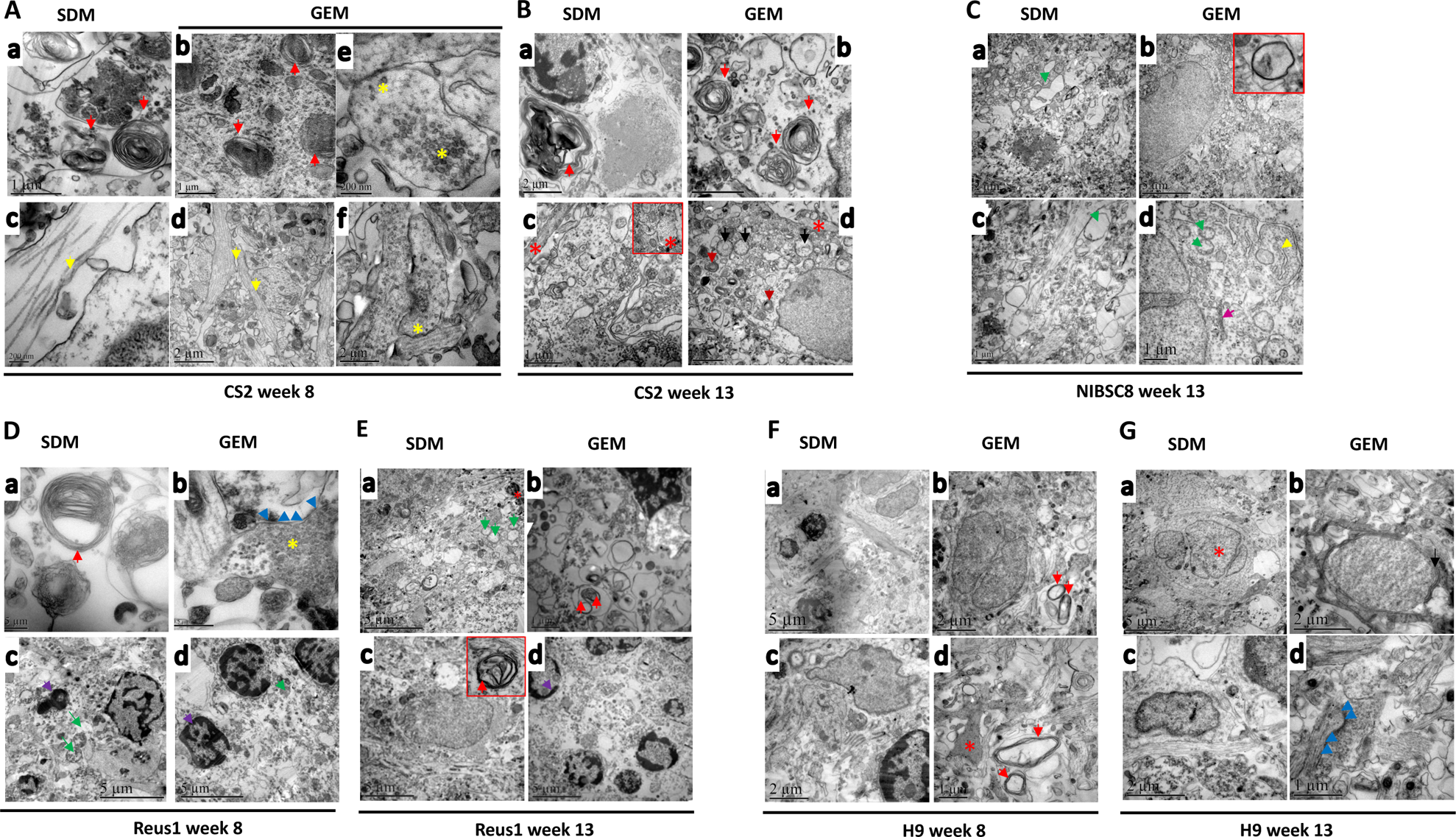
Electron microscopy of bMPS differentiated in SDM and GEM. (A). CS2 line at week 8 of differentiation shows presence of myelin (a, b red arrow), microtubule-rich processes (c,d yellow arrow), synaptic terminals (e,f, yellow asterisk) and microtubule-rich dendrites. (B) CS2 line at week 13 of differentiation shows presence of myelin (a,b red arrows, microtubule-rich processes (c,d red asterisks), and presence of edematous mitochondrial profiles with altered structure (d, black arrows). (C) NIBSC8 cell line differentiated for 13 weeks shows swollen mitochondria (a,c d, green arrows). GEM samples showed better organization and cytoarchitecture preservation, and cells with abundant cytoplasm rich in organelles, such as Golgi (d, pink arrow) and reticulum (d, yellow arrow). The insert shows a fiber in the myelination process. (D) Reus1 line at week 8 of differentiation shows presence of myelin (a, red arrow), synaptic terminal with synaptic vesicles (b, yellow asterisk), postsynaptic density (b, blue arrows), apoptotic bodies, pyknotic nucleus (c,d purple arrow), and mitochondrial profile with some structural anomalies (c,d green arrow) were evident. (E). Reus1 line at week 13 of differentiation shows presence of edematous mitochondrial profile (a, green arrow) and apoptotic bodies, pyknotic nucleus (d, purple arrow). The presence of myelin can also be observed (b, c red arrows). (F) H9 line differentiated for 8 week shows a vast network of extensions in the neuropile, myelin and myelination processes (b, red arrow panel) which were next to more electron-dense and compact cytoplasmic extensions (d, red asterisk), which resembles oligodendrocyte extensions. (G) H9 line at week 13 of differentiation shows multinucleated cells (a, red asterisk); GEM evidenced more synaptic contacts (d, blue arrows) and the myelinated process (b, black arrow).

### 3.5 Ca^2+^ transient differences between SDM and GEM

Finally, by measuring calcium signaling, we found that bMPS cultured in both media conditions changed the Ca^2+^ flux patterns during differentiation. Calcium transients were measured using fluo-4 which is a single-wavelength fluorescent Ca^2+^ indicator. Change in fluorescence over resting fluorescence intensity (delta F/F) was quantified (Supplemental Figure 6A and B). Interestingly, the delta F/F plots show that the week 15 SDM bMPS closely resembles the oscillation waveform of week 10 GEM bMPS, with sharp spikes suggesting a faster rise time (Supplemental Figure 6A-C). In contrast, for the GEM bMPS at week 15, their average peak amplitude drastically decreased, and their oscillation waveform exhibited a “plateau-like” structure. From the delta F/F plots, the average peak amplitude and interburst interval were calculated and compared between SDM and GEM cultures over time. The average peak amplitude was comparable between SDM and GEM cultures at week 10, but was drastically different between SDM and GEM cultures at week 15 with a significantly decreased average peak amplitude in GEM cultures (Supplemental Figure 6D-E). There were no statically significant differences in interburst interval, with a trend suggesting an increase in interburst interval in the week 10 and week 15 GEM cultures compared to SDM cultures. Interestingly, the interburst interval of week 15 SDM cultures and week 10 GEM cultures were comparable to similar mean values (Supplemental Figure 6D-E).

This point aligns with the Z-projections for the week 10 SDM and GEM organoids as well (Supplemental Figure 6C top). In addition, the Z-projections show more calcium staining along dendrites and axons in the week 15 SDM compared to the week 15 GEM bMPS, correlating to the delta F/F plots as well (Supplemental Figure 6C bottom).

Overall, when comparing week 10 conditions to week 15 conditions, the week 10 GEM bMPS show similar patterns to the week 15 SDM bMPS. Their oscillation waveform, average peak amplitude, interburst interval, and Z-Stack projections are similar in nature. This information combined suggests that GEM organoids are more mature at week 10 compared to the SDM organoids at week 10. Then at week 15, while the SDM organoids seem to mimic the activity of GEM organoids at week 10, week 15 GEM organoids showed decreased overall activity. Further analysis is necessary to understand whether this change is based on decreasing functionality, fewer synapses at week 15 versus week 10 or a plateau being reached.

These results suggest that GEM cultures are electro-chemically active and respond properly to depolarizations and the ensuing calcium influx response.

## 4. Discussion

Brain organoids represent a significant leap forward in microphysiological systems (MPS) to model human disease and screen novel drugs in a safe manner. Brain MPS recapitulate several key developmental aspects including genetic signatures of the brain development (38,39).

However, neurons still represent the majority of their cellular composition, with glial cells being substantially underpopulated (2,40). In our studies, using two iPSC and two ESC lines, we present the optimization of a system that enhanced astrocyte and oligodendrocyte populations. We demonstrated via immunofluorescence and bioluminescence assays that GEM significantly increases O4, MBP and GFAP expression. Genetic analysis confirmed a significant increased expression of *GFAP* and an upward trend of *MBP* expression in GEM treated cultures. Analysis of TEM images suggested differences between SDM and GEM, with potentially enhanced myelination, synaptic vesicles, and synaptic terminals in GEM cultures, though further analysis is needed. Finally, we also demonstrated through structural and functional experiments that GEM supports neurite outgrowth and cell migration, and that GEM cultures are electro-chemically active, mature faster, and respond properly to depolarizations and the ensuing calcium influx response.

An advantage of GEM over commercial media is that the key factors driving and enhancing the glial populations are known and can be easily modified to tailor culture needs. We previously showed that Neurobasal Plus medium in combination with B27 plus supplement also increased the glial populations in comparison to the Electrophysiology kit (23), however an advantage of GEM is that cultures can be modulated in a more flexible manner than when proprietary formulations are used.

In this improved model, neurons, oligodendrocytes, and astrocytes arise in the same system, allowing for a complex multi-cellular crosstalk that better reflects *in vivo* development.

Astrocytes and oligodendrocytes crosstalk via gap junctions, exchanging metabolites and signaling molecules (41–43). The physiological implications of this crosstalk influence metabolism and neuronal survival, auto and paracrine regulation of factors that promote synaptogenesis, OPC maturation, and myelination (44–46), and expansion of astrocytic networks that ultimately help shape neuronal circuitry and network activity (47). In our studies we observed that GEM-derived astrocytes better recapitulated the morphology of human primary culture astrocytes, appearing to induce a fibrous phenotype. Fibrous astrocytes are characterized by long-fibrous processes localized throughout the white matter in the CNS (9). While no particular marker has been associated with selective molecular signatures that could distinguish astrocytic phenotypes, the intermediate filament protein, glial fibrillary acidic protein (GFAP) is a canonical marker used to identify astrocytes throughout the CNS (48). Aligned with our results, other groups have found that iPSC-derived astrocytes resemble human morphology (49,50). The structural and functional plasticity of neuronal networks is partly governed by astrocytes, and as astrocytes age, they lose their ability to modulate and promote neurite outgrowth (35–37). In our studies we observed GEM significantly enhanced the neurite outgrowth and cell migration at week 13 of differentiation, supporting the cellular and molecular environment for both neuronal and astrocytic networks.

Weaved into the brain circuitry with neurons and astrocytes, myelinating oligodendrocytes play an array of functions. Beyond rapid propagation and fidelity of electrical transmission, myelinating oligodendrocytes also have roles in information processing (51). They can modulate the release of neurotransmitter at presynaptic terminals and can form functional synapses with neurons (14,52). Thus, oligodendrocytes have a prime and active role in modeling neuronal networks.

The importance of cellular composition and cellular cross talk between astrocytes and oligodendrocytes is evidenced in neurological diseases such as Alexander disease (AxD), where the loss of astrocytes leads to demyelination and oligodendrocyte death (41), ultimately affecting neuronal connectivity and loss of synapses (20). Emerging evidence has shed light on astrocytes increasingly becoming implicated in a variety of human diseases from leukodystrophies and congenital epilepsy syndromes, to neurodegenerative diseases such as ALS, Huntington’s and Alzheimer’s diseases (53). Given the significant enrichment of astrocytes and oligodendrocytes in our system, we consider our model a useful tool for precision medicine, where patient-derived bMPS can be created to study the pathophysiology of disease and screen for drugs that can better inform safer regenerative therapies.

## Funding

This study was supported by NIH grants T32-ES007141-38 and K12-GM123914.

## Author contributions

Conceptualization LS and IEMP. IEMP designed the study and performed most of the research. LS and IEMP wrote the manuscript. LD provided experimental support and neurite outgrowth quantification. PL and SAM provided TEM imaging. CR provided experimental advice and created and validated the NPCs bank utilized in the research. DAE provided calcium imaging analysis, XC and DJZ designed and provided the sec Nanoluc ESC lines. All authors contributed to the article and approved the submitted version.

## Acknowledgments

We acknowledge the Integrated Imaging Center at Johns Hopkins University. We thank Wilmer Core Grant, Microscopy module, EY001765. We thank the Institute of Biology of Fluminense Federal University for allowing access to its transmission electron microscope (JEM 1011) facility from PMEIB/UFF.

## Conflict of Interest

Standard organoids (SDM) are licensed to AxoSim, New Orleans, LA, United States. LS consults AxoSim. The rest of the authors declare no conflict of interest.

## Abbreviations

BMPS: Brain microphysiological systems
BDNF: Brain Derived Neurotrophic Factor
ESC: Embryonic stem cells
GDNF: Glia Cell Derived Neurotrophic Factor
GEM: Glia-enrichment medium
GFAP: Glial fibrillary acidic protein
IGF-1: Insulin like growth factor 1
iPSC: Induced pluripotent stem cells
MBP: Myelin Basic Protein
NPC: Neuroprogenitor cells
NF-H: Neurofilament Heavy chain
O4: Anti-sulfatide antibody O4
PLP1: Proteolipid protein 1
PDGFRa: Platelet-derived growth factor receptor alpha
PDGF-AA: Platelet-derived growth factor AA
SDM: Standard differentiation medium
T3: 3,3,5-Triiodo-l-thyronine

## Supplemental Figure Legends

**Supplemental 1.**
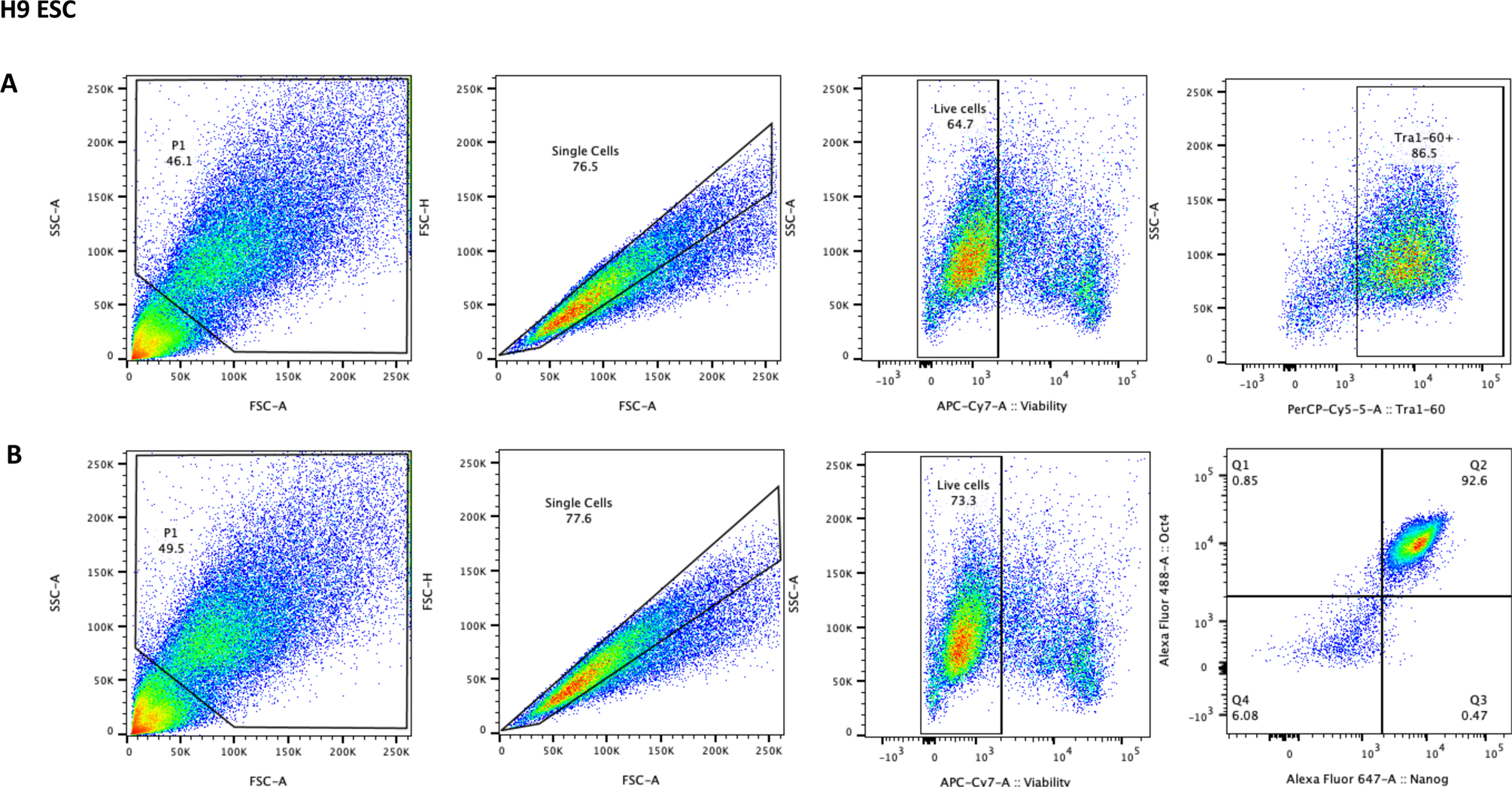
Flow cytometry characterization of human ESC line H9. Representative flow cytometry plots showing the gate strategy to identify live cells expressing the canonical pluripotency stem cell markers TRA-1-60 (A) and Oct 4 and Nanog (B).

**Supplemental 2.**
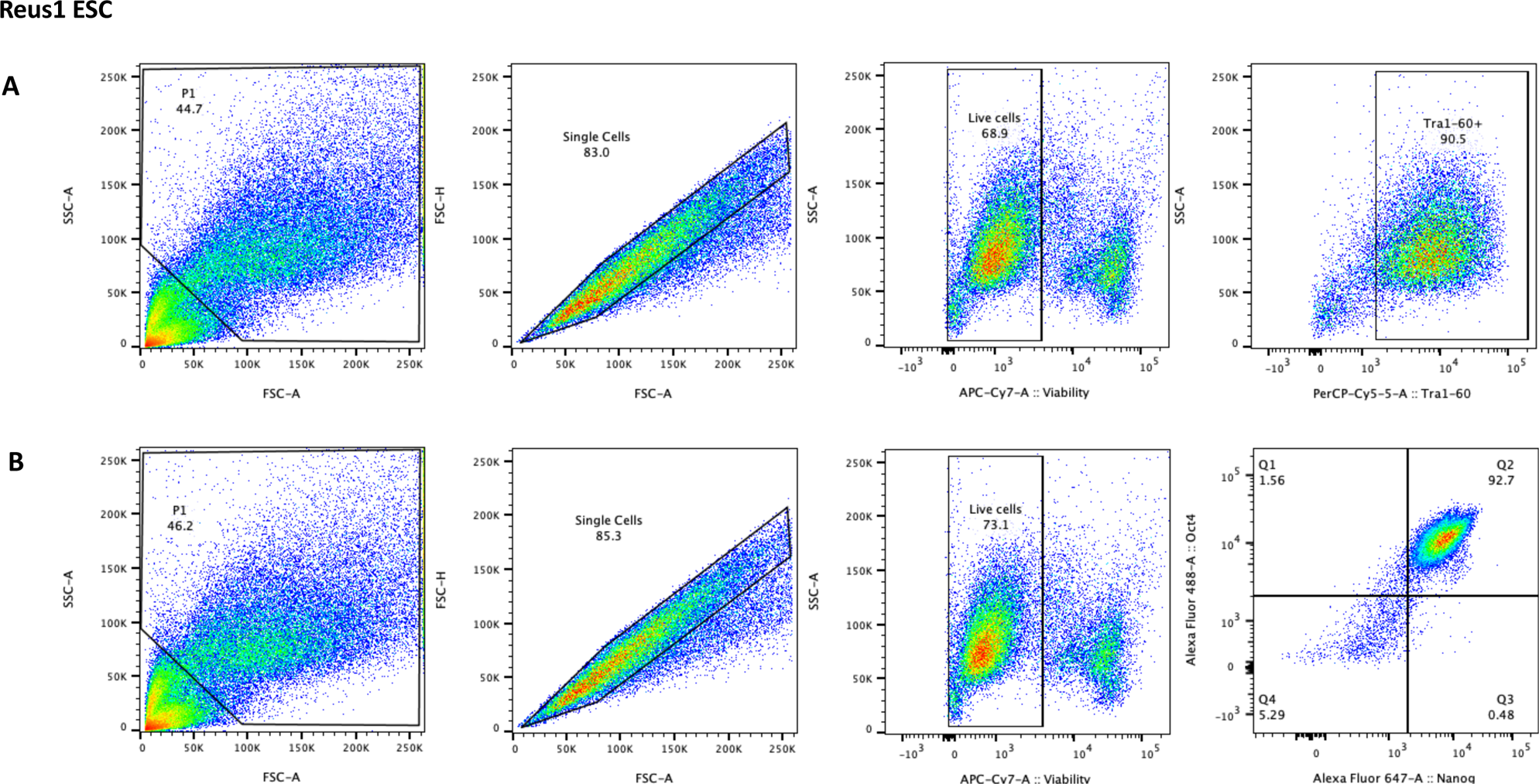
Flow cytometry characterization of human ESC line Reus 1. Representative flow cytometry plots showing the gate strategy to identify live cells expressing the canonical pluripotency stem cell markers TRA-1-60 (A) and Oct 4 and Nanog (B).

**Supplemental 3.**
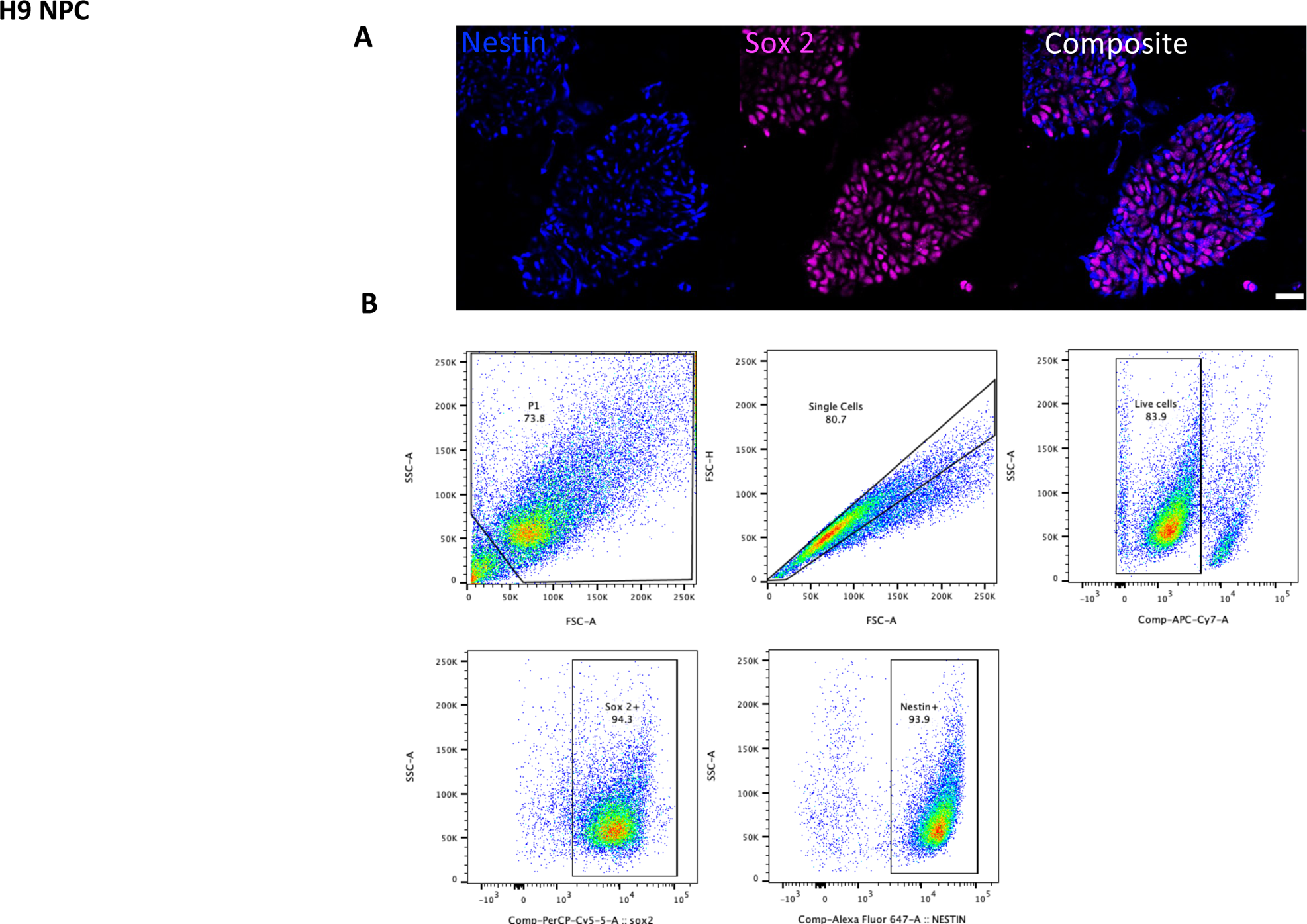
Immunofluorescence and flow cytometry characterization of H9 NPCs. (A) NPCs are labeled with antibody against Nestin (blue), and Sox 2 (magenta) markers. The scale bar represents 50 µm. (B) representative flow cytometry plots showing the gate strategy to identify live cells expressing the NPC markers Sox 2 and Nestin.

**Supplemental 4.**
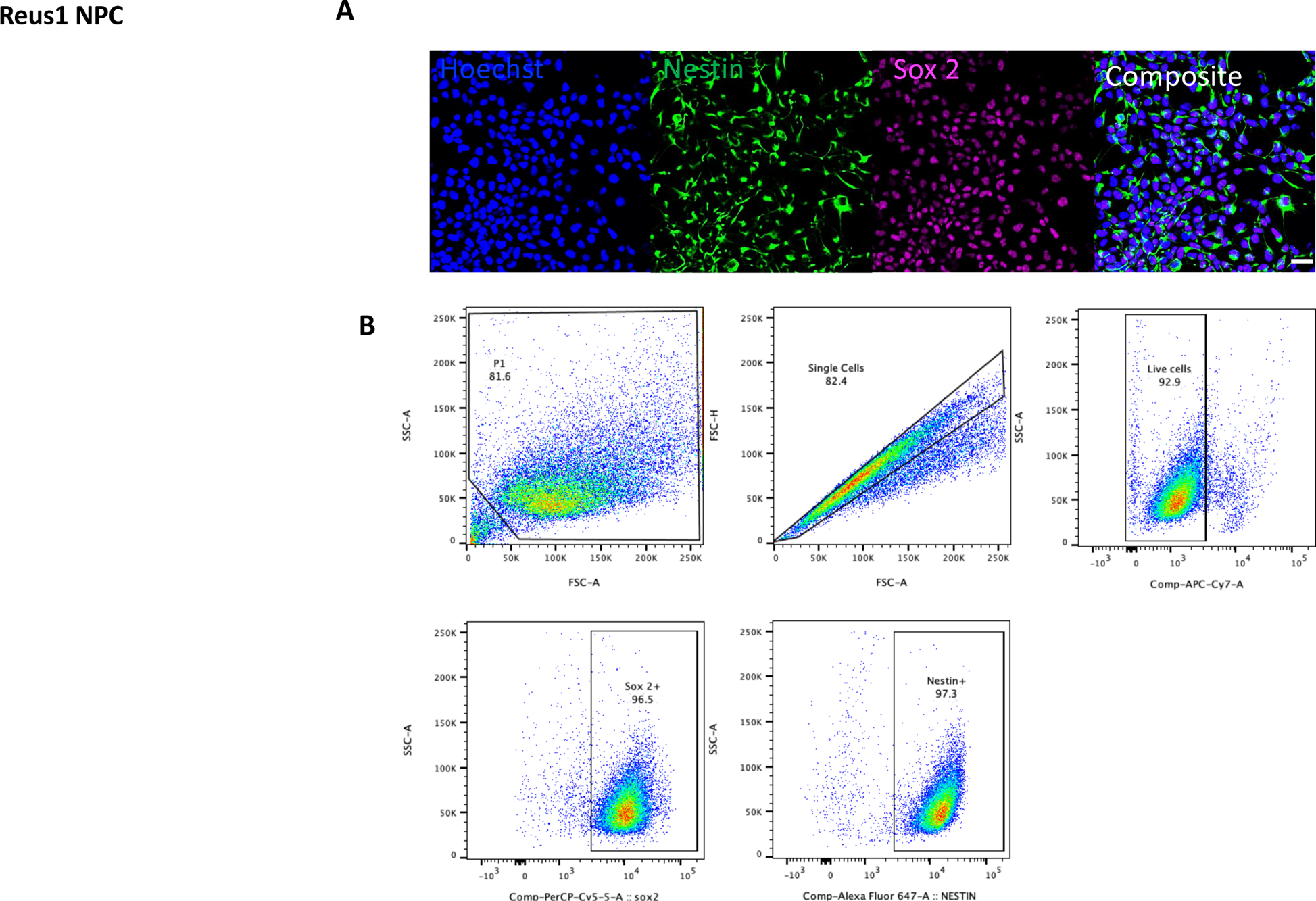
Immunofluorescence and flow cytometry characterization of Reus 1 NPCs. (A) NPCs are labeled with antibody against Nestin (green), and Sox 2(magenta) markers. The scalebar represents 50 µm. (B) representative flow cytometry plots showing the gate strategy to identify live cells expressing the NPC markers Sox 2 and Nestin.

**Supplemental 5.**
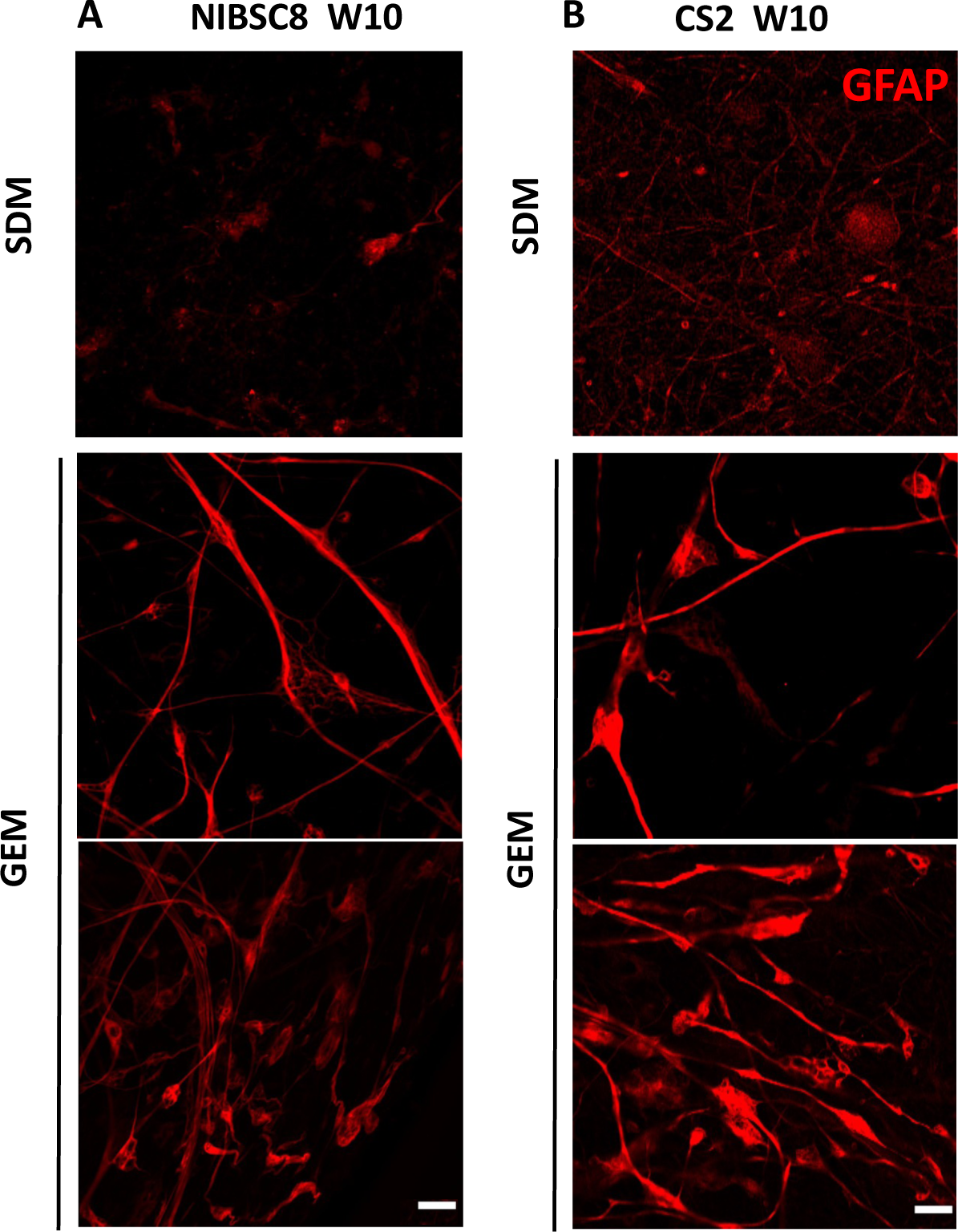
GEM astrocytes closely recapitulate morphology of human primary cultures. Representative images from at least three independent experiments per cell line show astrocyte morphology in SDM (top panel) versus GEM (bottom panel) conditions at week 10 of differentiation of iPSC-derived bMPS from NISBC8 (A) and CS2 (B) lines. The scale bar represents 50 µm.

**Supplemental 6.**
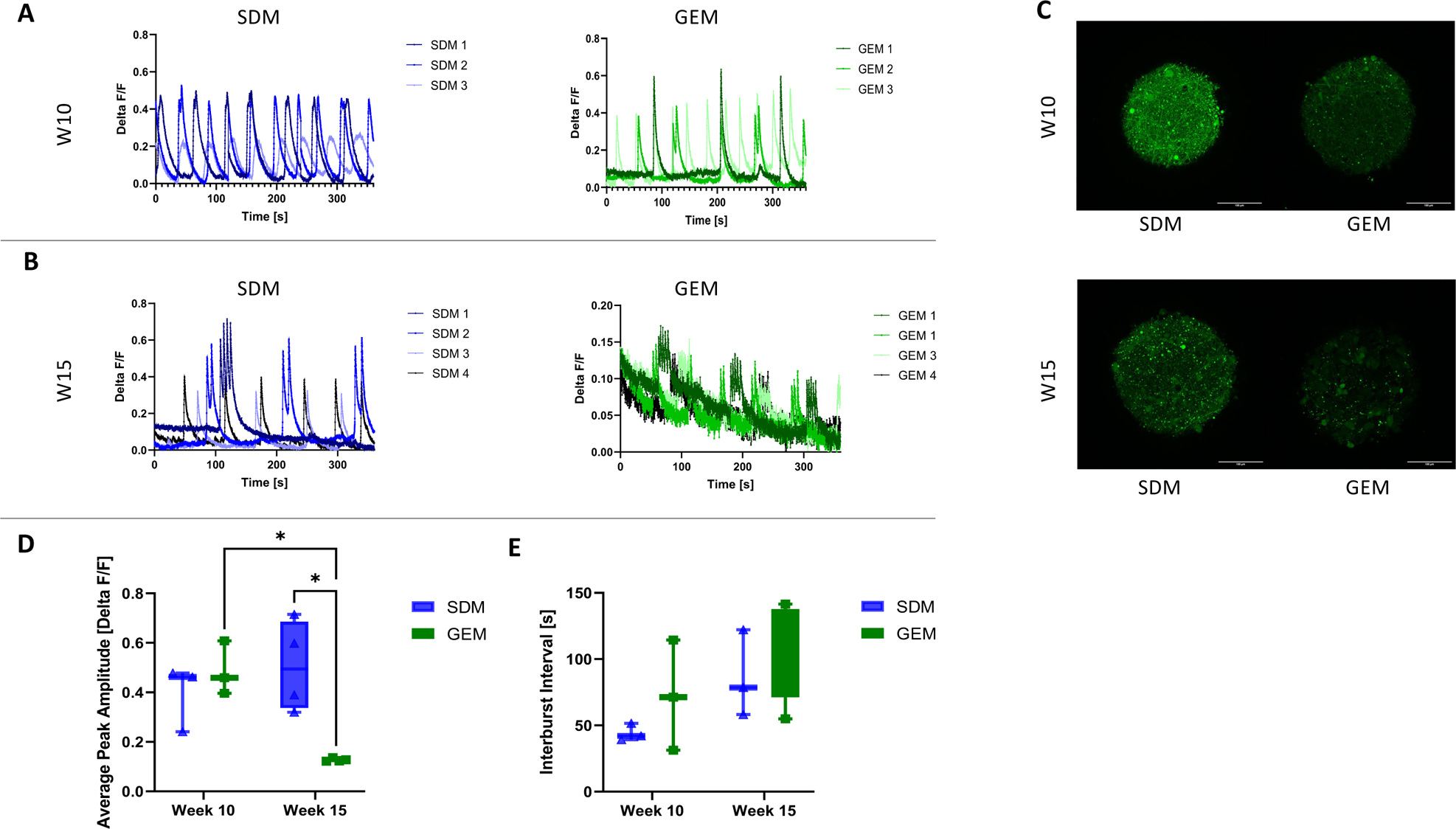
Calcium signaling analysis of SDM and GEM bMPS. The change in fluorescence over resting fluorescence intensity (delta F/F) in SDM and GEM bMPS at week 10 (A) and week 15 (B) of differentiation, accompanied by representative images of calcium staining (C). Three to four bMPS were used per condition. Comparison of the average peak amplitude (D) and interburst interval (E) between time of differentiation and medium composition. The scale bar represents 100 µm. A two-way ANOVA test with Holm-Sidak post-hoc test was used for statistical analysis and p < .05 was considered significant.

